# Potential acoustic signatures of stress in black soldier fly (*Hermetia illucens;* Diptera: Stratiomyidae) larvae

**DOI:** 10.64898/2026.03.06.709542

**Authors:** CD Perl, O Escott, G Reiss, A Crump, M Barrett

## Abstract

Black soldier fly larvae (BSFL) have quickly become one of the most farmed animals in the world. However, little is known about how to monitor stress and welfare in these animals. The difficulty of welfare assessment is compounded by the fact that BSFL live in their feed and prefer darkness. This behaviour makes it challenging to observe potential welfare indicators without inducing stress via disturbing the larvae or moving them into the light. However, acoustic devices may be able to pick up signatures of stress in the population even while they are out of sight, allowing for remote monitoring of animals in natural conditions (in the feed and/or in the dark). Acoustic monitoring of this type has been deployed for the detection of insects in stored grains, suggesting this method holds some promise for assessing insect behavioural signatures. In this study, we aimed to identify general, acoustic signatures of stress in BSFL by recording them during exposure to two stressors (light or shaking) or in a low-stress control condition. Our data suggest there are consistent differences in the acoustic recordings of the ‘non-stressed’ and ‘stressed’ conditions that may indicate the animals’ behaviours shift consistently in response to stress. Ultimately, the data suggest acoustic monitoring may hold promise for larval behaviour and/or welfare assessment and should be further explored in response to a variety of stressors across the larval life stage.

## Introduction

Passive acoustic monitoring is a minimally invasive method for assessing the welfare of vertebrate livestock, with animals’ sounds (or lack thereof) revealing their behaviour and/or emotional states (Ramos Niño *et al*., 2025). For instance, prolonged, high-frequency calls can indicate distress in piglets during weaning when separated from their mothers (Weary *et al*., 1999). Chicken pecking sounds can indicate successful feed intake and growth, while vocalisations can also signal thermal discomfort or issues with air quality (Soster *et al*., 2025). The ‘murmuring’ sound of cattle while ruminating can be monitored via audio recordings, as it has a significantly lower frequency than calls made during other activities (Meen *et al*., 2015). Passive acoustic monitoring has also proved effective for measuring feeding activity in invertebrate livestock, particularly in species of *Penaeus* shrimp (Smith & Tabrett, 2013; Peixoto & Soares, 2025; Zhang *et al*., 2025). New artificial intelligence technologies are even being explored to improve vocalization analysis, and thereby behaviour and welfare monitoring (Herlin *et al*., 2021; Manikandan & Neethirajan, 2025).

Insect mini-livestock, such as the black soldier fly (*Hermetia illucens*, Diptera: Stratiomyidae), are the basis of a novel animal agriculture industry, focused on providing animal feed to support food security for a growing human population (van Huis, 2013; United Nations, 2022). Nearly 5 trillion insect larvae are expected to be slaughtered as food and feed annually by 2033 (with the vast majority being black soldier fly larvae; BSFL). This makes insect agriculture the fastest-growing and largest animal agriculture industry on the planet (by number of individual animals; McKay & Shah, 2025). Despite a relative lack of knowledge about insect sentience at the juvenile lifestage (Gibbons *et al*., 2022), precautionary ethics, economic incentives, and industry best practice documents all emphasise the importance of considering farmed BSFL welfare (while further research into sentience is conducted; Birch, 2017; IPIFF, 2019; Barrett & Adcock, 2023).

Accordingly, tools are needed to assess BSFL behaviours and putative welfare states (Barrett & Fischer, 2023). BSFL are especially challenging to monitor, as they live in dense aggregations in their feeding substrate, completely hiding them from view and making it difficult to follow individual animals. Between 2,000 and 14,000 BSFL may live in a single bin of feed (or more, in trough systems), and they tend to cluster within the feed as they eat (Shishkov *et al*., 2019). Consistently bringing larvae to the surface for visually monitoring behaviour and welfare can cause stress via both disturbance/handling and exposure to light (Barrett *et al*., 2023a; Cattaneo *et al*., 2025; Durosaro *et al*., 2025). Methods of assessing larval welfare without disturbance and in the dark will be essential to non-invasively assess welfare.

Acoustic monitoring of larval behaviour and welfare may solve these challenges. While no studies have attempted acoustic monitoring with BSFL to date, this method has long been used with other insects hidden from view. Stored grain pests are often detected using acoustic monitoring, which catches the sounds of feeding activity or motion (Shuman *et al*., 1997; Mankin *et al*., 2021). In conservation settings, acoustic monitoring has also been regularly employed for detecting insects (Riede & Balakrishnan, 2025). For instance, freshwater insect populations can be monitored using acoustic signals (Desjonquères *et al*., 2024). Thus, BSFL behaviours and/or welfare states might also be acoustically assessed.

Here, we use full-spectrum audio loggers to explore the potential for acoustic monitoring in BSFL welfare and production. We assess: 1) if acoustic monitoring devices can detect BSFL activity while they are in their substrate; and 2) if stressors (disturbance, light) produce consistent acoustic signatures when compared to lower-stress rearing conditions.

## Materials and Methods

### Insect Source and Husbandry

Five-day-old larvae were supplied by AgriGrub (Wisbech, UK). Fifteen plastic trays (60 x 40 x 20 cm) were filled with 9kg of feedstock each and seeded with 10,000 BSFL (gravimetrically determined). Trays were placed randomly onto racking units within a 12 m containerised insect unit at the University of Leeds Research Farm. Sample sizes (for trays) were based on the number of trays that could be successfully reared and recorded from simultaneously, limited by space and equipment availability. The number of individuals per tray was chosen with the aim of replicating industrial conditions as closely as possible, thus with at least 10,000 BSFL per tray. The unit was set to 27 °C and 60% relative humidity.

### Recordings under Higher and Lower Stress

Recordings were taken of 15 and 16 day old BSFL, once feedstock was dry enough, as a result of bioconversion, that larvae could be manually sieved from the substrate (60cm diameter, 125 mm mesh aperture size; Russell Finex, Feltham, UK). This was done to improve the ability to detect acoustic signals compared to in-substrate recordings based on our pilot test. On day 15, larvae were sieved from the substrate in half of the trays, and combined as a group. The remaining half of the trays were sieved and assigned to treatments the following day (day 16) and then subjected to the same procedure.

On day 15 and 16 of rearing, 200 larvae were subsampled from the aggregated population and used to determine average larval weight with an analytical balance accurate to 0.001 mg. On both days, 200 larvae weighed 37g (0.185g per larva). Using this figure, 2000 larvae were gravimetrically portioned into each of nine experimental bins (18,000 larvae per day). Three trays were randomly assigned to each treatment (Control, Physical Disturbance, Light Exposure), and an additional tenth tray remained free of larvae (Empty treatment) to serve as a control for room noise (e.g., HVAC systems, human movement). All ten trays were recorded continuously for six hours using full-spectrum audio loggers (AudioMoths v.1.2.0, Open Acoustic Devices, UK). These were in a waterproof casing and suspended inside each tray, angled with the microphone toward the bottom where the larvae were. Trays were not sound-proofed during the experiment, due to our pilot trials demonstrating this did not improve sound quality.

‘Physical Disturbance’ trays were manually shaken for 1 minute every 1 hour during the recording period, and left in the dark. ‘Light Exposure’ trays were left untouched but had two 1000 lumen LED lights (JCB Work Lights, Stratford-Upon-Avon, UK) mounted above each tray. Control trays were left in the darkness and not disturbed for six hours. Following the six-hour recording period, larvae used for the experimental trays were killed by freezing. To avoid acclimatization, larvae were only used once for acoustic monitoring. Experimenters were not blinded to treatment during data collection or analysis.

### Recording and Statistical Analysis

First, recordings were extracted and imported into Audacity v 3.7.7 Audio Editor (https://www.audacityteam.org/) for listening and visual inspection.

To quantify differences across the conditions, two acoustic metrics were calculated: 1) root mean square energy (RMS), which measures the energy or loudness of a signal, often referred to as ‘power’; and 2) acoustic complexity index (ACI), which measures temporal variability of sound intensity in a signal and should capture any repetitive, high-frequency, transient signals produced by larval activity.

For each day, 20 ten-minute segments were sampled at random from each condition after the first disturbance/stressor period (or a matched period in controls where there was no stressor applied). Therefore, 60 segments were obtained for each condition (3 trays x 20) and 20 segments from the reference/empty tray on each day. The two metrics were then calculated on each segment in Python (v.3.14). ACI was calculated using the sci-kit maad library (Ulloa *et al*., 2021) and RMS Power was calculated using the librosa library (McFee *et al*., 2015).

All statistical analyses were conducted in R (version 4.5.1). Treatment differences in ACI and RMS power were analysed using linear mixed effects models constructed using the package ‘lme4’ (Bates *et al*., 2015). Minimum adequate models, specifically to ascertain the most suitable random effect structure, were selected via a partial F-test and comparison of AIC and BIC scores. In the instance of a significant difference between models, the model with the lowest AIC and BIC score was selected. Model comparison indicated that ‘tray’ nested in ‘experimental day’ and modelled as a random intercept provided the best fit for mixed effects models. Treatment was modelled as a fixed effect. Significance of fixed effect model terms was ascertained via the ‘lmerTest’ package (Kuznetsova *et al*., 2017), and post-hoc pairwise comparisons between treatment levels was conducted via Tukey tests from the package ‘emmeans’ (Lenth, 2024). For all tests, alpha was set to 0.05, model assumptions were assessed by plotting residuals against fitted values.

### Ethics Statement

Although no legal protections exist for insects used in research, we still aimed to minimise any pain and distress experienced by insects where possible. First, we conducted a pilot test to inform the likelihood of success of our experimental paradigm, given there are no prior studies on acoustic recording of BSFL. Next, we aimed to choose acute but non-lethal stressors that minimized the risk of nociception and tissue damage while still causing fitness-relevant stress exposure.

Larvae were killed via freezing at the end of the study. We recognize that this is not recommended as a humane method for terrestrial invertebrate euthanasia without prior anesthesia (Leary *et al*., 2020; Bakker *et al*., 2026; Fischer *et al*., 2026). However, this was the only method available that complied with institutional safety policies, and it is still generally considered to be the most feasible, safe, and widely-practiced option for mass killing research insects.

## Results

### Acoustic signatures of BSFL activity

In all recordings, including the empty bin without larvae, the dominant signal was the HVAC system, which appears as a static noisy signal below 1.5 kHz. However, in the empty bins on both days, there were no transient signals at higher frequencies (10 - 20 kHz); these signals were only observed in the bins containing larvae (Figure 1), and represent the sounds of larvae rapidly moving against each other. Therefore, acoustic monitoring can detect larval movement while out of substrate.

**Figure 1.**
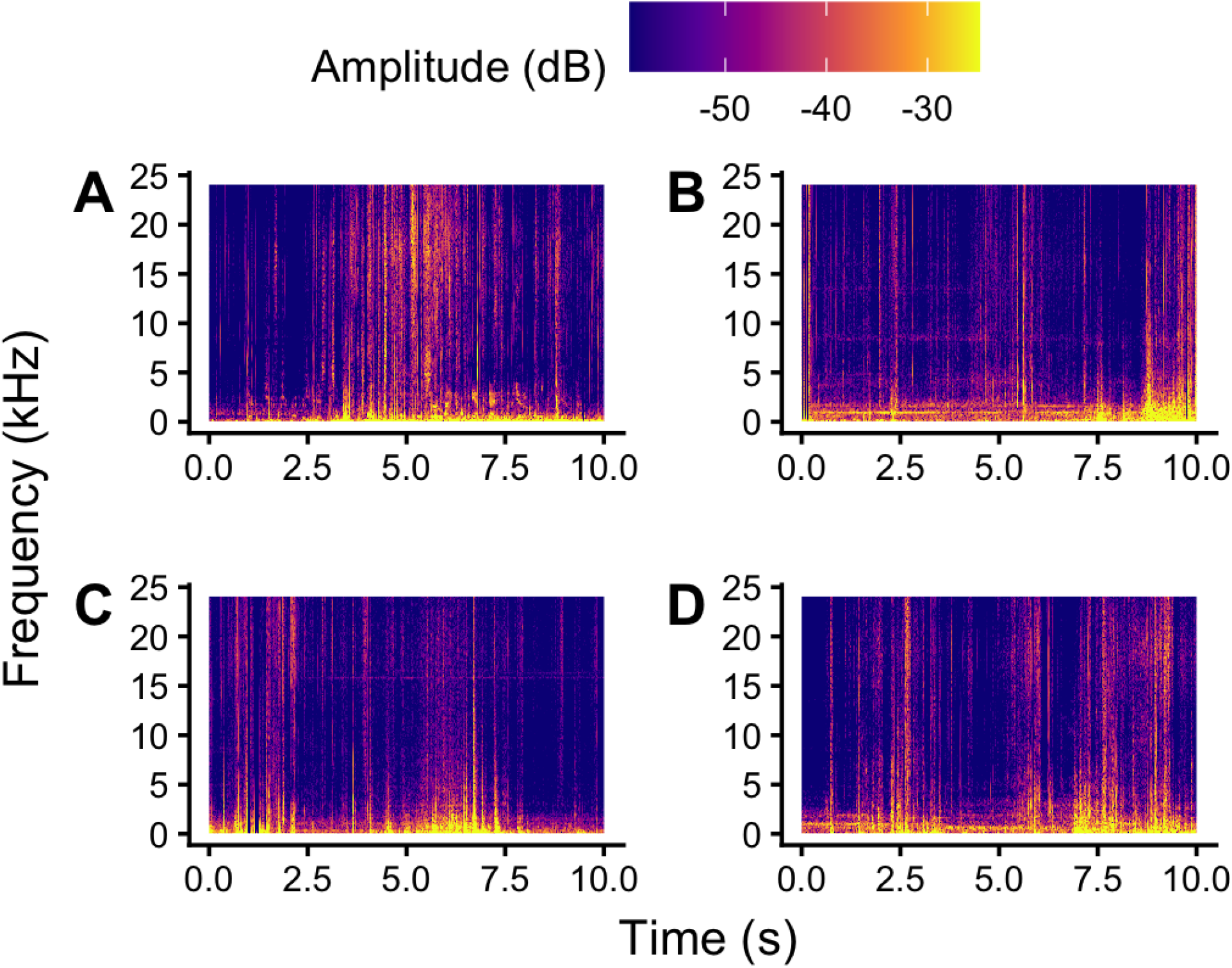
Example spectrograms. **A)** Control tray of larvae (no physical disturbance, in the dark); **B)** Physical disturbance treatment; **C)** Light exposure treatment; and **D)** Empty tray without any larvae. Larger versions of these spectrograms can be found in Figure S1. The RMS analysis shows a statistically significant difference (Figure 2B) between trays with larvae and the empty trays, further highlighting the fact that larval activity can be assessed with acoustic signals even with relatively loud background (HVAC) noise in the system.

### Differences between stressed and non-stressed treatments

There were significant differences in ACI between the treatments (Figure 2A, LMM; F_3,400_ = 44.52, p < 0.001). The Control treatment produced a higher ACI score than the Light Exposure (t_139,400_ = 7.50, p < 0.0001) or the Empty treatment (t_182,400_ = 10.12, p < 0.0001). There was no significant difference in the ACI produced by the Control and the Disturbed treatments (t_139,400_ = 1.92, p = 0.22). The Disturbed treatment produced a higher ACI score than either the Light (t_139,400_ = 5.58. p < 0.0001) or the Empty treatments (t_182,400_ = 8.33, p < 0.0001). The Light treatment produced a higher ACI score than the Empty treatment (t_182,400_ = 3.12, p < 0.02).

**Figure 2.**
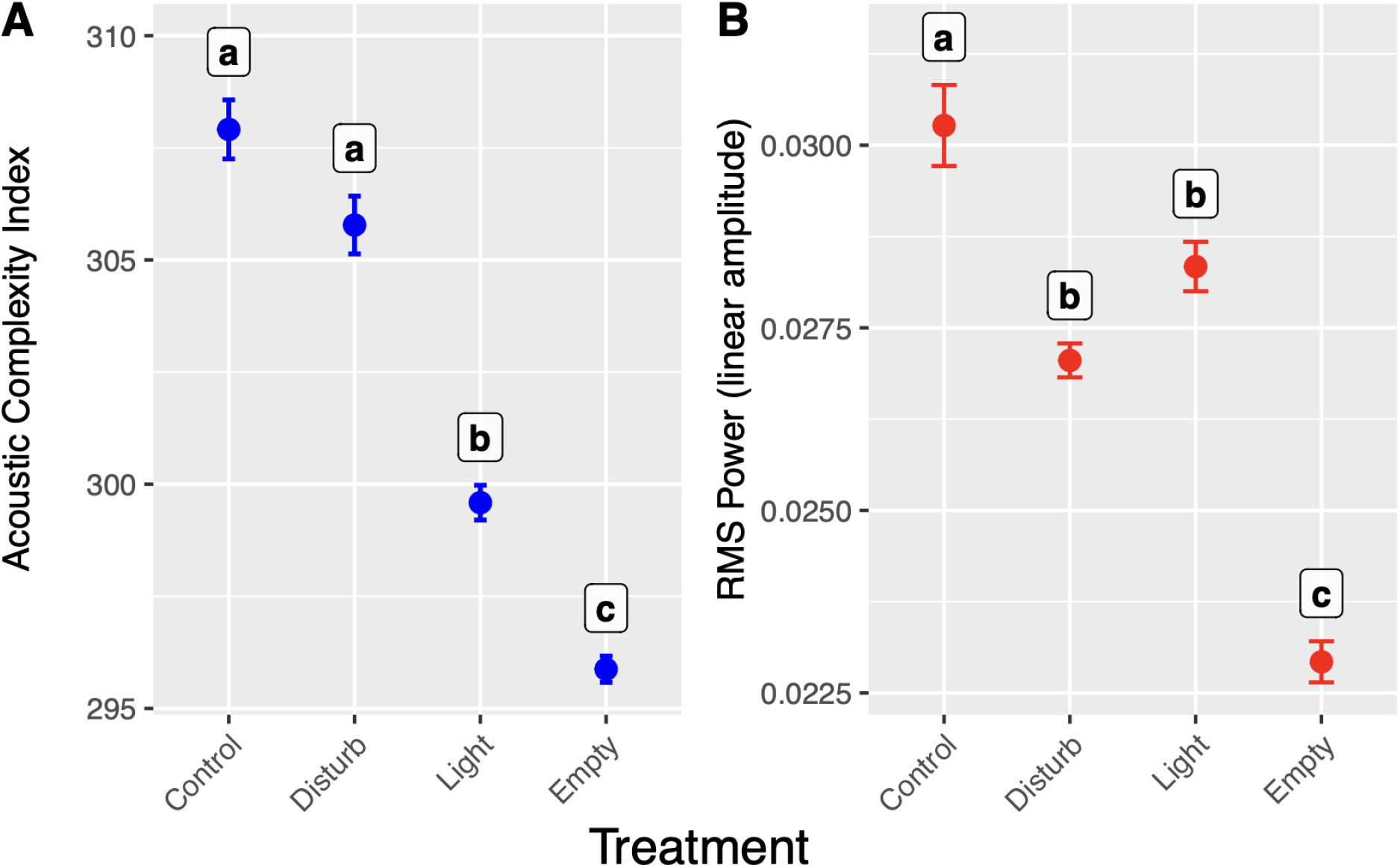
Acoustic welfare indicators across treatments. **A)** The acoustic complexity index is highest in trays experiencing minimal stress (Control) and those exposed to Physical Disturbance, while Light Exposure trays had reduced ACI. (B) The root mean square (RMS) power produced by the trays was highest in the Control treatment and lower in both stressed treatments (Physical Disturbance and Light Exposure). Empty trays without larvae had the lowest ACI and power. Letters indicate statistically significant differences among treatments (p < 0.05).

There were significant differences in the power produced between the treatments (Figure 2B, LMM; F_3,400_ = 29.82, p < 0.001). Control treatments produced higher power than Disturbed treatments (t_136,400_ = 4.43, p < 0.001), Light treatments (t_136,400_ = 2.66, p < 0.05) and Empty treatments (t =_193,400_ = 9.24, p < 0.0001). Disturbed treatments produced a higher power than Empty treatments (t_193,400_ = 5.20, p < 0.0001) but were not significantly different from Light treatments (t_136,400_ = 1.77, p = 0.29). Light treatments produced higher power than Empty treatments (t_193,400_ = 6.82, p < 0.0001).

## Discussion

We found that acoustic monitoring should be explored as a promising indicator of behaviour and welfare in BSFL. Signatures of larval activity (transient, high-frequency sounds) could be reliably distinguished from background noises (like HVAC systems) that are common in intensive insect production facilities (which the University of Leeds Research Farm was built to model). This supports the idea that acoustic monitoring should be further explored for assessing BSFL activity and welfare on farms. The BSFL mostly produced sounds in the 15-20kHz range. These sounds putatively reflect activity patterns as BSFL are not known to communicate via sounds (though some other insect larvae do: Low *et al*., 2021). Importantly, our recordings occurred when larvae were out of substrate (as may happen on farms for 24-72 hrs while being fasted before slaughter or, on some farms, in preparation for pupation; reviewed in Barrett et al. 2023). Further tests of acoustic monitoring of actively feeding BSFL in substrate would be valuable to assess this monitoring tool across the entire larval lifestage.

Our data demonstrated that exposing BSFL to two different stressors (Physical Disturbance or Light Exposure) impacted acoustic cues when compared to less-stressed controls. Light Exposure significantly reduced both the acoustic complexity index (ACI) and power, while Physical Disturbance significantly reduced power. ACI indicates the complexity of captured sound, with higher scores reflecting more irregular and diverse frequencies. While continuous light exposure caused a loss of acoustic complexity consistent with reports of larval photophobia (e.g., Cattaneo et al. 2025), the stress of intermittent physical disturbance did not result in a similar loss, with this treatment statistically indistinguishable from less-stressed controls.

While this finding may mean that ACI is not a uniform indicator of poor welfare, the inconsistent results could also represent a difference in the magnitude and timing of stressor application in our study design. The light stressor was applied continuously throughout the six-hour recording, while the physical disturbance stressor was applied for only 1 minute of every sixty minutes throughout the recording (to avoid the sound of manual shaking interfering with the acoustic monitoring). If handling were applied continuously, as was light, this may influence ACI. This seems especially plausible as, although the physical disturbance treatment was not statistically distinguishable from controls, it did trend in the same direction as the light exposure treatment. Nevertheless, our ACI data indicates that constant light-associated stress may cause a change in behaviour that reduces energy across some frequencies non-uniformly, whereas our intermittent physical disturbance treatment did not result in the same pattern. Ultimately, these data suggest that reductions in sound intensity (e.g., power) could indicate poor larval welfare under standard farming conditions, as both stressors reduced sound intensity.

Lower power is, putatively, a sign of reduced larval activity. While both of our stressors were purposefully sublethal and acute, lethal or chronic stressors might result in different activity patterns, and thereby variation in the degree or direction of any ACI and power changes. For example, lethal overheating is a common welfare concern for BSFL (Barrett et al. 2023, and see Schow-Madsen *et al*., 2025 for a discussion of sublethal heat exposure impacts). Heat may increase insect activity, as they are poikilotherms (having a metabolic rate largely determined by environmental temperature); further, lethal heat causes BSFL to attempt to escape their rearing trays (Fuhrmann *et al*., 2025), which could increase larval activity patterns on recordings. Alternatively, a different lethal stressor, like disease, could reduce activity due to lethargy, which is a common symptom of insect diseases, including in BSFL (She *et al*., 2023). Ultimately, testing these acoustic cues for novel stressors will be necessary to understand how generalizable acoustic signaling may be for assessing BSFL welfare. Different stressors may also generate different frequencies (though we were unable to confirm this with our data due to limited sample sizes); the variable responses of ACI to different stressors lends some support to this hypothesis.

Our BSFL findings strongly indicate that other commonly-reared insect species may also benefit from acoustic monitoring. For instance, both adult crickets and black soldier flies use sound for mating behaviour (Giunti *et al*., 2018; Rowe *et al*., 2024). Yellow mealworms, also reared en masse and without many on-farm welfare assessment tools (Barrett *et al*., 2023b), are common grain pests, so recent research has developed sensors to detect their feeding activity (Kadyrov *et al*., 2024). These tools could also be adapted for passive acoustic behaviour and welfare monitoring.

In conclusion, passive acoustic monitoring is a low-cost, minimally-invasive tool that can successfully indicate BSFL responses to stress during out-of-substrate periods in industrial rearing conditions. Sound intensity is a good candidate for a general indicator of at least acute, sublethal stress in BSFL. ACI deserves further research as it responded to only one stressor in our study, though this could reflect the variability in stressor application. Our data serve as a proof-of-concept to verify passive acoustic monitoring for farmed insect welfare assessment. Nevertheless, further investigation is needed across different farmed species and the stressors they are likely to encounter.

### Animal Welfare Implications Statement

BSFL are among the most-farmed animals in the world, but methods for assessing their welfare, or even monitoring changes in activity patterns, remain underexplored. This paper provides a passive on-farm method for recording larval activity and reliably distinguishing it from background noise. Moreover, the technology reliably distinguished the acoustic cues of BSFL housed in lower-stress v. higher-stress conditions (at least, for the acute stressors of light and disturbance).

## Acknowledgements

We thank AgriSound (particularly David Geary) for providing audiomoths and helping with data analysis and study design. We thank the technician team at the University of Leeds research farm for assistance gathering data and rearing animals.

## Supplemental Materials

**Figure S1.**
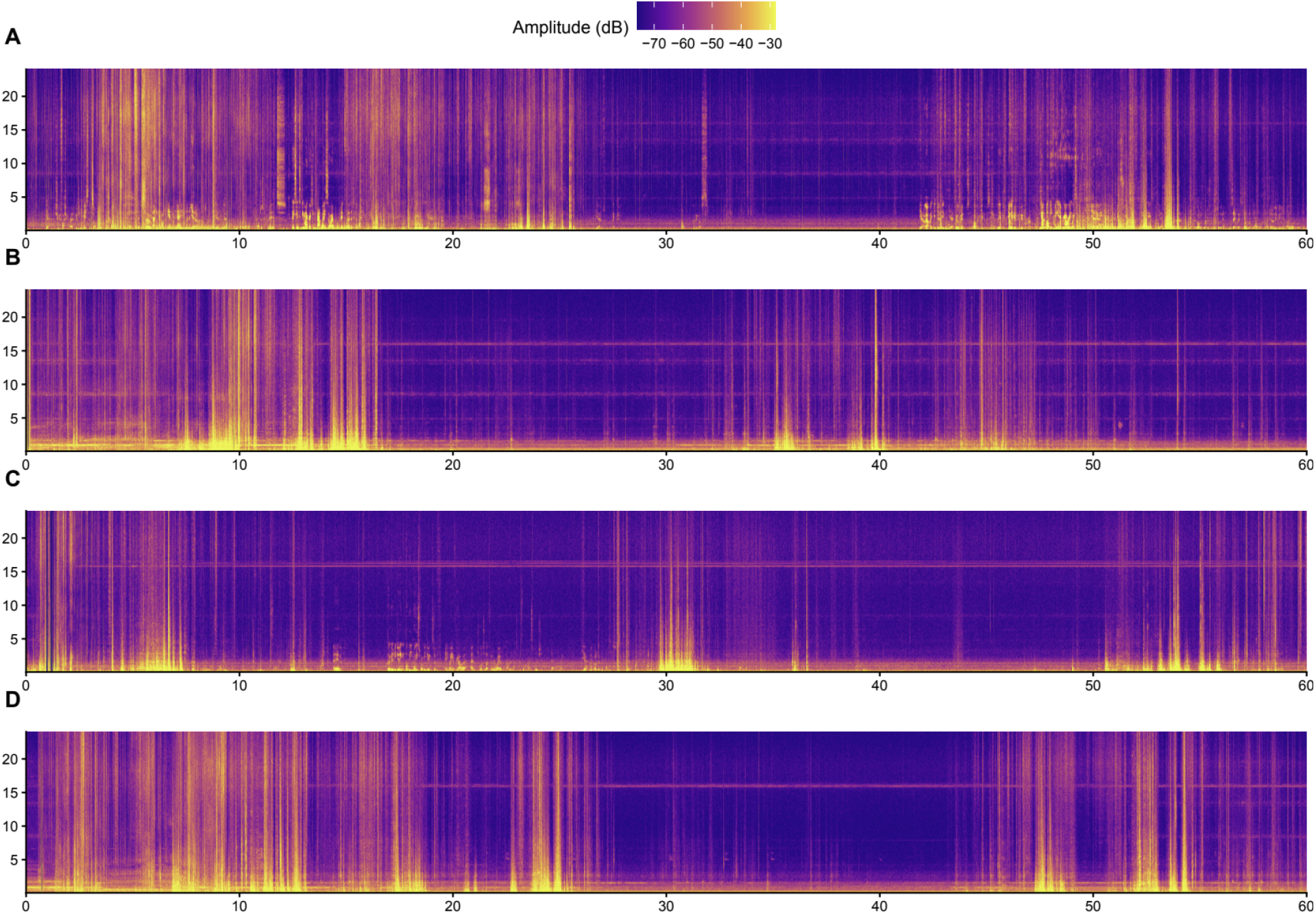
Larger versions of the example spectrograms from Figure 1. **A)** Control tray of larvae (no physical disturbance, in the dark); **B)** Physical disturbance treatment; **C)** Light exposure treatment; and **D)** Empty tray without any larvae.

